# Theoretical framework and experimental validation of multiplexed analyte quantification using cross-reactive affinity reagents

**DOI:** 10.1101/2023.11.24.568623

**Authors:** Sharon S. Newman, Linus A. Hein, Alexandra M. Adams, H. Tom Soh

## Abstract

Gold standard immunoassays depend on specific affinity reagents for accurate molecular quantification. Any cross-reactivity of affinity reagents, wherein the reagent non-specifically binds to unintended molecules, can create false positive binding signals and result in inaccurate quantification of analytes. Mitigating cross-reactivity represents one of the greatest challenges in molecular diagnostics, and remains an unsolved problem. To instead overcome the effects of cross-reactivity, we present a mathematical framework that uses generalized binding equations and noise estimation to enable the use of multiple cross-reactive reagents for multiplexed molecular quantification. As a proof-of-concept, we experimentally demonstrate accurate quantification of a small molecule for which no specific affinity reagents are available, even at high concentrations of a cross-reactive molecule. Furthermore, this robust schema yields well-defined bounds of quantification that make it easier to assess the quality of assay results and predicts under which conditions assay performance is likely to break down. This work turns cross-reactive affinity reagents, which were previously a liability, into an asset for achieving accurate quantification of analytes.

---

The ability to measure medically relevant analytes with high accuracy is crucial as their concentrations often provide valuable insights into disease status and therapeutic response. To achieve accurate molecular quantification, gold standard immunoassays are highly reliant on specific affinity reagents. If an affinity reagent exhibits cross-reactivity, wherein the reagent binds to unintended molecules, false positive binding signals can be produced and thus hinder the accurate quantification of protein and small molecule analytes.

Despite this reliance on specificity, even antibodies, which are a benchmark for affinity reagents, suffer from cross-reactivity. This is evidenced by a large scale study of 11,000 antibodies, in which over 95% exhibited cross-reactive binding, often to non-target analytes that were in high abundance or shared sequence homology [1]. A more recent study of 153 antibodies observed that 84% were cross-reactive, and notably, about 47% of these unintended analytes bound even stronger than the intended target protein itself, posing a substantial obstacle to accurate quantification [2]. Unfortunately, researchers often incorrectly assume that reagents are specific and thus inadvertently dismiss cross-reactive signals as constant background signals, leading to inaccurate conclusions [3]. Cross-reactivity thus poses a serious obstacle for diagnostic accuracy and sensitivity — a problem that is only exacerbated as multiplexed immunoassays scale up [4].

Several methods have been developed to mitigate the effects of cross-reactivity through both assay design and affinity reagent development methods. For example, gold-standard methods such as enzyme-linked immunosorbent assays (ELISAs) rely on the recognition of different epitopes by multiple affinity reagents to drastically lower the effects of cross-reactivity [3, 4, 5]. Unfortunately, smaller analytes, such as short peptides (e.g., hormones) and small molecules (e.g., drugs and metabolites), often lack multiple epitopes that can be recognized simultaneously, and thus depend on single-reagent assays such as competitive ELISAs. For these assays, the accuracy of the readout is entirely dependent on the availability of high-specificity reagents. However, developing such reagents is extremely challenging, especially for low-molecular-weight analytes, and remains an unsolved problem for many important biomarkers [6, 7, 8]. As such, there is an urgent unmet need for strategies that can achieve accurate analyte quantification even if the affinity reagents themselves inevitably retain some level of cross-reactivity.

As a solution to the cross-reactivity problem, we have developed a mathematical framework for accurate quantification that is robust to the cross-reactivity of affinity reagents — which is, to our knowledge, the first such approach described in literature. To overcome the challenges of using cross-reactive affinity reagents, our framework comprises three features: 1) A generalized equilibrium-based model [9, 10] to account for cross-reactivity, 2) Employment of multiple affinity reagents to resolve individual contributions of analytes, and 3) Accounting for noise to ensure robust measurement and solution stability. We present experimental results to validate our approach with a fluorescence-based assay by quantifying a small molecule metabolite, kynurenine (kyn), for which no specific affinity reagent is available. Using cross-reactive affinity reagents, we quantified kyn in solution with moderate to high concentrations of cross-reactive non-target interferent. As expected, a conventional model for analyte-reagent binding that does not account for cross-reactivity provided highly inaccurate results with deceptively narrow ranges of quantification. Conversely, using our framework we accurately determine a narrow concentration range for kyn despite even higher concentrations of cross-reactive interferent. Furthermore, we present how our framework can predict the quantitative precision of such cross-reactive assays with visual intuition, which makes it possible to both identify scenarios in which such assays can achieve robust quantitation and identify opportunities for further optimization of assay performance. Our work thus offers an avenue to achieving robust analyte quantification even with cross-reactive affinity reagents, allowing for enhanced molecular measurements without relying on stringent affinity reagent selection or complex assay design.

## Results and Discussion

### A model for cross-reactive affinity reagents

To begin, we provide a model that describes the total signal output *s*_*i*_ from an affinity reagent *A*_*i*_ in a solution of *n* analytes. When cross-reactive, *A*_*i*_ can bind to any one of the *n* analytes *T*_*j*_ at a time, where *j* ranges from 1 to *n*. Each of these *n* interactions occur with distinct association constants 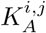, and can be depicted as follows:

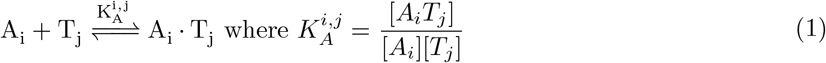

Using the basic assumptions stated in Methods 1.1, we can model the equilibrium state of this system and derive the following equation for the normalized output signal *s*_*i*_ from reagent *A*_*i*_:

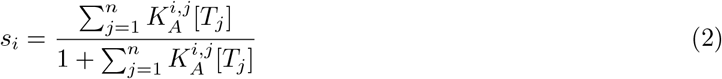

For simplicity, we analyze and visualize this model (and our framework) in the context of a cross-reactive affinity reagent *A*_1_ binding to just two analytes: a desired analyte *T*_1_, and an interferent analyte *T*_2_ (Fig. 1a). In this case, the output signal *s*_1_ from reagent *A*_1_ reduces to:

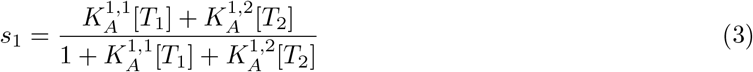

**Figure 1.**
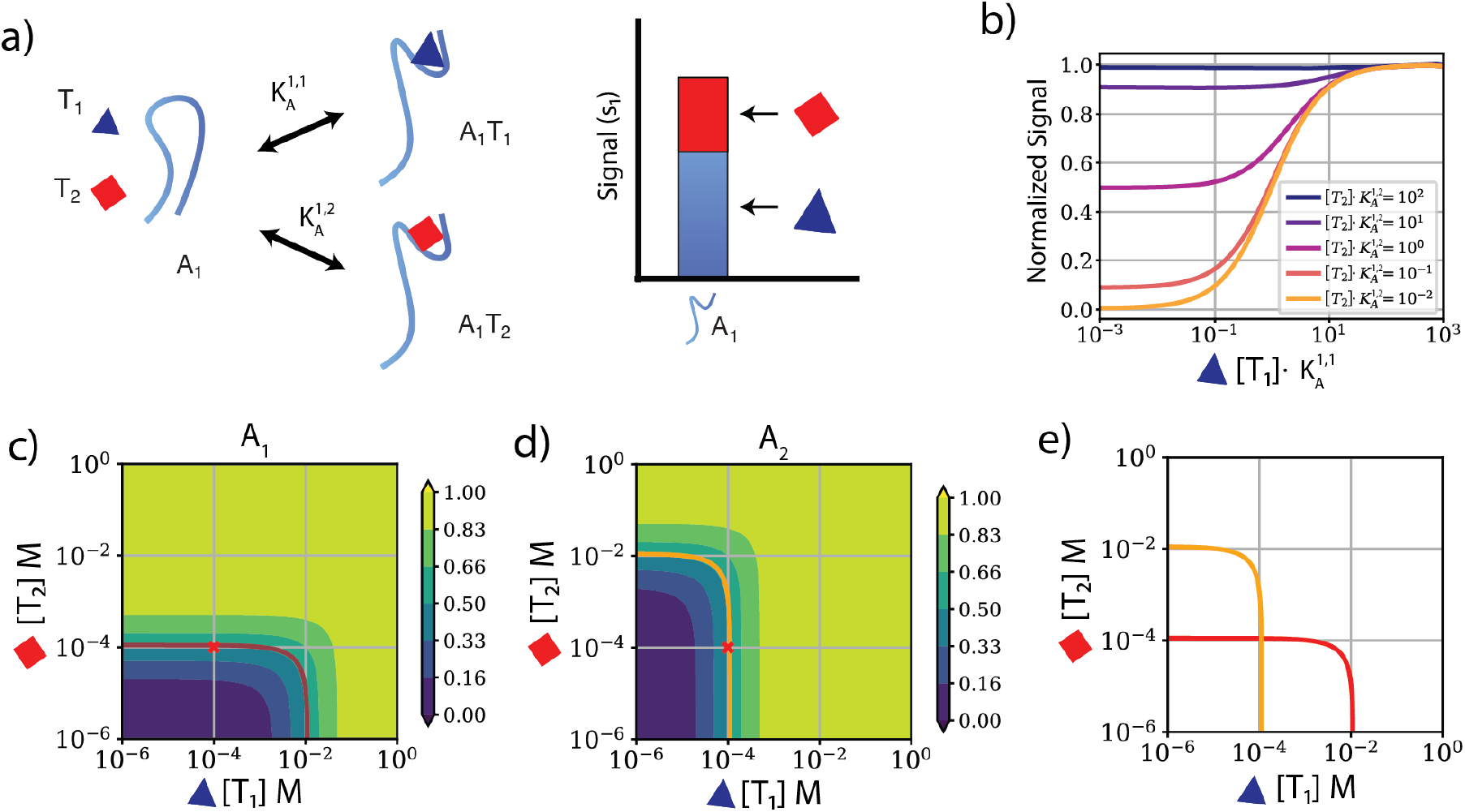
The signal output from a cross-reactive affinity reagent is greatly affected by the concentration of non-target analyte. **a)** An affinity reagent, *A*_1_ (aptamer in this cartoon) can bind to both the target analyte (*T*_1_) and any cross-reactive analyte (*T*_2_). When incubated with a mixture of *T*_1_ and *T*_2_, *A*_1_ produces a signal *s*_1_ that is composed of contributions from both *T*_1_ and *T*_2_. **b)** The dynamic range of the signal intensity *s*_1_ from reagent *A*_1_ decreases with increasing [*T*_2_]. Colors represent different values of [*T*_2_] relative to the reagent’s association constant, 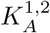. **c**,**d)** Heatmap of signal across the feasible range of [*T*_1_] and [*T*_2_] for **c)** reagent *A*_1_, with 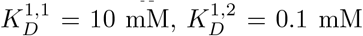 and **d)** reagent *A*_2_ with 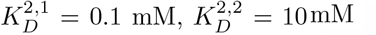. The color scale represents the normalized signal *s*_1_ and *s*_2_ from each affinity reagent, where blue is no signal (= 0) and yellow is saturated binding (= 1). The red cross indicates an example concentration where both [*T*_1_] and [*T*_2_] are 0.1 mM. Red and orange curves in b and c respectively illustrate the range of possible concentrations that could produce the same output signals. **e)** The overlap of these curves is the solution for [*T*_1_] and [*T*_2_] given the indicated values of *s*_1_ and *s*_2_ from *A*_1_ and *A*_2_, respectively.

Since *A*_1_ can bind to both *T*_1_ and *T*_2_, *s*_1_ is a function of contributions from [*T*_1_] and [*T*_2_] (Fig. 1a). In Figure 1b, we plot Equation 3 for [*T*_1_] at different concentrations of *T*_2_(which is also a function of the association constants 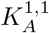 and 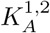). As [*T*_2_] increases, the lower limit of the [*T*_1_] to *s*_1_ binding curve shifts upward, i.e., the magnitude of *s*_1_’s response to an increase in [*T*_1_] also depends on [*T*_2_]. This shift in the lower limit of the signal highlights the cause for concern if [*T*_2_] changes or is not expected to affect *s*_1_, as in the case of assuming specificity with the conventional Langmuir isotherm (when [*T*_2_] = 0) [11].

To further elucidate the effect of different combinations of [*T*_1_] and [*T*_2_] on the signal *s*_1_ from *A*_1_, we have plotted Equation 3 as a two-dimensional heatmap for a scenario in which our affinity reagent binds to *T*_1_ and *T*_2_ with equilibrium dissociation constants *K*_*D*_ = 1*/K*_*A*_ of 10 mM and 0.1 mM, respectively (Fig. 1c). Given only the value of *s*_1_, we cannot accurately determine a value for [*T*_1_] or [*T*_2_], as Equation 3 yields a wide set of solutions that form the red highlighted L-curve shown in Figure 1c. In this example, [*T*_1_] could range from 0 to 10 mM.

To narrow the possible space of concentrations for *T*_1_ and *T*_2_, we can introduce another affinity reagent *A*_2_ to provide an additional signal *s*_2_. To this end, these two affinity reagents need to be distinct in terms of having linearly independent association constants for the two analytes (i.e., 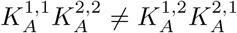). The signal from each affinity reagent can be handled separately, assuming the affinity reagent concentrations are much lower than the analyte concentrations. Since the *K*_*A*_ values in our example are linearly independent, the solution space for *A*_2_ is different from that of *A*_1_ (Fig. 1d). By using both affinity reagents to measure the same mixture of two analytes, we obtain two signals that correspond to the L-curves highlighted on the two heatmaps. Superimposing these curves, the overlap point represents the space of [*T*_1_] and [*T*_2_] that simultaneously explains the signal *s*_1_ from *A*_1_ and the signal *s*_2_ from *A*_2_ (Fig. 1e). The concentration of the sample can now be resolved analytically to show that [*T*_1_] and [*T*_2_] in this example are both 0.1 mM.

This approach of overlapping signals from affinity reagents is theoretically applicable for an arbitrary number of affinity reagents and analytes using Equation 2^1^. However, as we increase the number of analytes and affinity reagents beyond two, the solution space from each analyte is unlikely to neatly overlap in a single point due to the compounded effects of noise, such that analytical solutions become intractable. This means that analytical solutions using Equation 2 are only dependable when the noise is negligible (see SI Section 2.2), and we must incorporate noise into our framework—which we do in the following section.

### Accounting for noise improves ability to find a solution for analytes

It is essential to account for the sample noise profiles associated with these signals to robustly resolve the analyte concentrations. To incorporate noise into our model, we derive the 95% confidence interval for the measurement signals. We use measurement replicates *s*_*i,r*_ to define upper and lower bounds on the mean signal 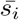, which contains the true noise-free signal *s*_*i*,true_ with a probability of 95%. We assume that our noise follows a Gaussian distribution with some standard deviation *σ*, such that (*s*_*i*,true_−*s*_*i,r*_) ∼ 𝒩 (0, *σ*^2^).

With *R* replicate measurements, we calculate the 95% confidence interval by first computing the mean signal intensity of our measured samples 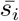:

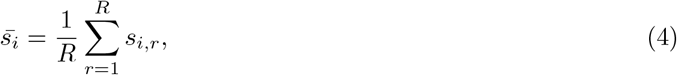

and the sample standard deviation *SD*_*i*_:

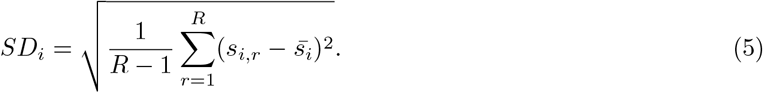

We can then denote the 95% confidence interval as:

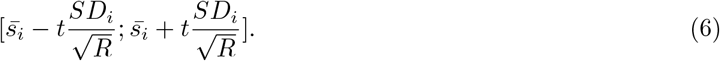

with *t* being the 97.5th percentile of the t-distribution with (*R* − 1) degrees of freedom. Defining 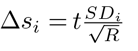, with 95% confidence we have:

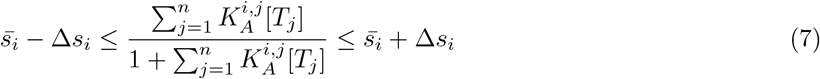

This confidence interval results in a thickening of the plotted curve, which encompasses the true analyte concentrations within a 95% confidence interval. The overlap of these broader curves for all the affinity reagents defines a set of analyte concentrations that, while not having any statistical guarantees, in practice, usually include the true analyte concentration while also accounting for measurement noise.

We can then solve for the upper and lower bounds of each analyte’s concentration range. In order to visualize concentrations spanning several orders of magnitude, we have been plotting this solution space on a log-log scale. However, we can more easily solve for these bounds mathematically in the linear domain. Thus we reformulate Equation 7 above into a constraint on a linear combination of the analyte concentrations [*T*_*j*_]:

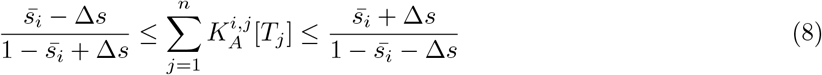

We can further constrain the size of the solution space by adding another set of inequalities that capture the physical limits of each molecule *T*_*j*_, which range from 0 to the solubility limit *T*_*max,j*_. These limits can also be further constrained by other physiological bounds. Based on these linear constraints, we can use off-the-shelf convex optimization solvers to calculate the range of concentrations expected for each analyte (Method 1.2).

### Experimental validation of our model to quantify analytes with a pair of cross-reactive affinity reagents

To experimentally verify our approach, we performed an assay to quantify the tryptophan metabolite kynurenine (kyn), for which no specific affinity reagent exists. We used affinity reagents by Yoshikawa and Wan et al. that were developed to specifically bind to many structurally-similar metabolites within the kynurenine pathway including xanthurenic acid (xa) [7], but none were found to be specific only to kyn. Specifically, we used a cross-reactive reagent (SK1) that binds to both kyn and xa, and a xa-specific affinity reagent (XA1). Employing the same fluorescence assay protocol they used, we quantified kyn in the presence of xa.

We first re-established the dissociation constants of the affinity reagents to both analytes. We confirmed that SK1 is indeed cross-reactive with a *K*_*D*_ of 0.13 mM for xa and ∼10-fold poorer affinity for kyn (*K*_*D*_ = 1.98 mM, Fig. 2a). We also determined that XA1 has a *K*_*D*_ of 0.64 and 264 mM for xa and kyn, respectively. We note that 264 mM is far beyond the solubility limit of kyn (19 mM in water [12]), and thus XA1 can be considered a highly selective affinity reagent. These values are comparable to those reported by Yoshikawa and Wan et al. (SI, Table 2-3).

**Figure 2.**
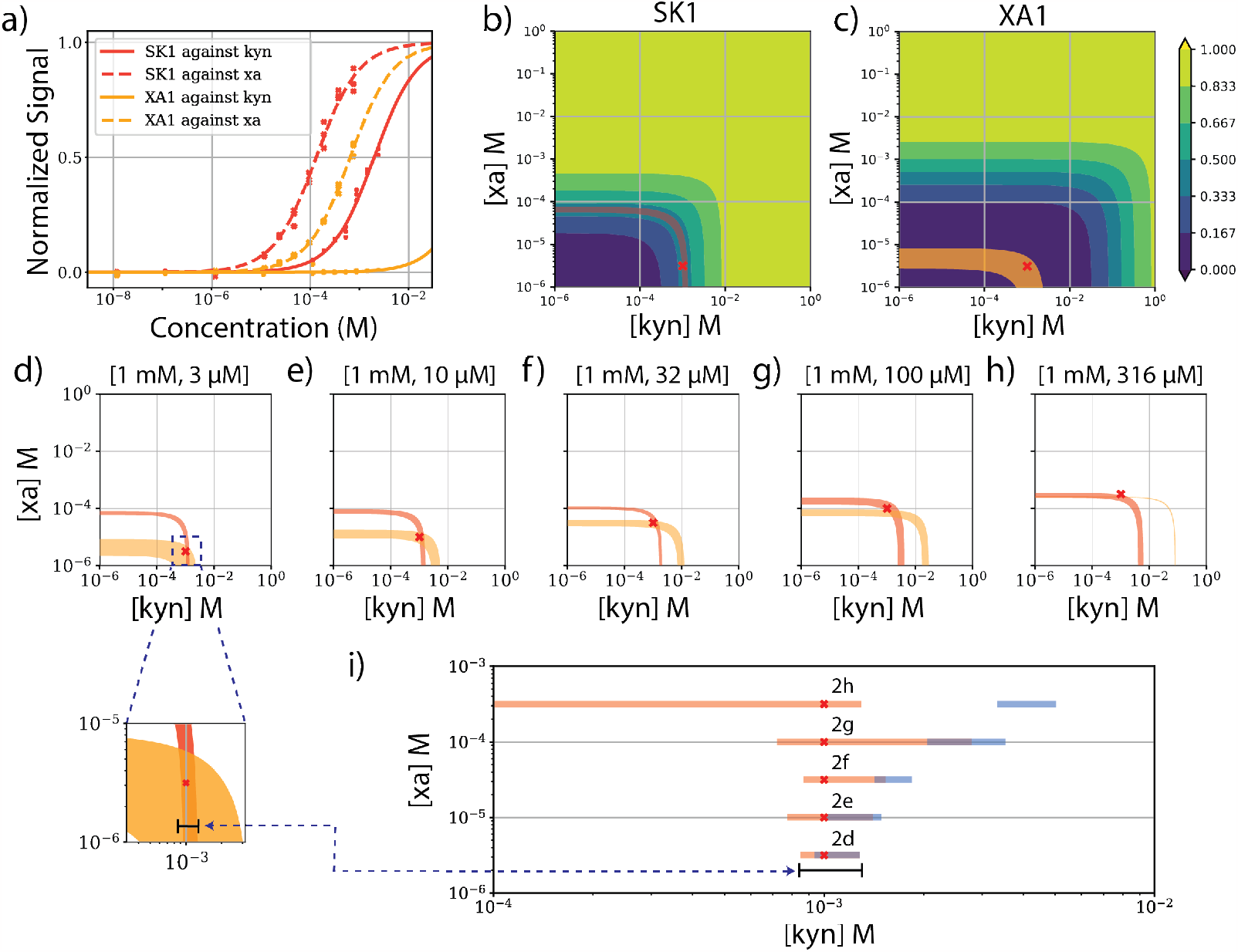
Experimentally assessing our quantification framework with cross-reactive affinity reagents. **a)** Normalized binding curves for affinity reagents SK1 (red) and XA1 (orange) to kynurenine (kyn; solid lines) and xanthurenic acid (xa; dashed lines). **b, c)** Signal heatmaps with feasible analyte concentrations from **b)** SK1 and **c)** XA1. Color scale shows normalized signal intensity. Highlighted L-curves (red and orange) indicate 95% confidence intervals for feasible analyte concentrations that could yield the normalized signal produced in the presence of 3 µM xa and 1 mM kyn (marked with a red cross). **d-h)** Overlapped 95% confidence interval curves produced by the two affinity reagents at the same kyn concentration (1 mM) but increasing xa concentrations. **d)** Zoomed view of overlapping region with bar indicating the estimated concentration range for kyn. **i)** Estimated concentration ranges of kyn for each sample mixture using our method (orange), compared to assumption of specificity (blue). Red crosses mark the true input concentration (1 mM kyn). Exact values presented in SI Table 4-5.

For initial demonstration of our capacity for kyn quantification in presence of xa, we created five samples. In each sample, the concentration of kyn was maintained at 1 mM, while the concentration of xa varied ranging from 3 µM to 316 µM. Based on the *K*_*D*_ fits, we can plot signal heatmaps for both SK1 (Fig. 2b) and XA1 (Fig. 2c), alongside the feasible concentrations from their respective signal readouts(Fig. 2d-h). We then convert these measurements into lower and upper bounds for the concentrations of kyn and xa using our above-described quantification method. As an exemplar, with an input concentration of 1 mM kyn and 3 µM xa, we obtain the signal as highlighted in Figure 2b-c. Using the signal from each affinity reagent by itself, we observe a wide range of possible kyn concentrations, spanning from 0-3 mM. By combining the information from both affinity reagents, as shown in Figure 2d, we greatly narrow down the feasible concentration of kyn. The true concentration of kyn (1 mM) falls within the calculated bounds of the overlapped curves (0.87–1.25 mM), which produces a narrow range with log error (= (log_10_(measured) −log_10_(true))) ranging from −0.06 to 0.10 (Fig. 2i).

To assess the performance of our model, we compared our results against a ‘naïve’ model, which does not account for cross-reactivity, by applying our confidence interval and linear programming framework to a basic Langmuir binding equation. The naïve model consistently overestimated the concentration of kyn in the presence of the interferent xa (Fig. 2i, blue; SI Table 4). In fact, the naïve model only predicted an accurate concentration range for kyn at the lowest concentration of xa. But even then, the range of feasible kyn concentrations was skewed towards higher values, with a log error range of −0.02 and 0.10. With increased [xa] in subsequent samples, the naïve model predicted narrow concentration ranges that are misleading and highly inaccurate since the lower bounds of these ranges consistently exceeded the true concentration. The naïve model’s quantitative prediction increasingly differed from reality with upper and lower bound log errors ranging from 0.01 to 0.69 log error. In contrast, by accounting for cross-reactivity, our model was able to calculate a range of kyn concentrations that were accurate even in the presence of extremely high [xa]. For the samples with xa concentrations below 100 µM, the calculated ranges were also very narrow with largest lower and upper log errors of −0.05 and 0.18 (Fig. 2i, orange; SI Table 5). For the samples with greater than 100 µM xa, these ranges became much wider, with the upper end of the log error extending out to 0.44 with no lower bound for the sample with 316 µM xa. Despite the wider ranges calculated for these high concentrations, it is important to note that our model predictions still reliably encompass the true concentration of kyn, in contrast to the fully inaccurate predictions of the naïve model. The wide quantification ranges provided for these higher xa concentrations are a result of fundamental limitations of the affinity reagents and assay design, and not of our framework^2^. We delve into predicting when such quantification ranges will be narrow, and how quantification can be improved in the following section.

### A model for predicting the range of quantification for assays

Our experimental quantification above demonstrated narrow quantification ranges within a certain regime of concentrations given the affinity reagents. Beyond such a regime however, the ranges become much wider and thus become less useful for conventional quantification use cases, despite being technically accurate. Here, to better understand what cases will produces narrow or wide ranges of quantification, we predict the ‘range of quantitation’ (ROQ), which is a metric for how tightly constrained the overlap of the confidence intervals is. We furthermore demonstrate how we can use such ROQ predictions to rationally improve quantification.

The ROQ will increase as an affinity reagent approaches saturation or when the signal output from the analyte is close to the assay’s background. These boundaries are highly dependent on the coefficient of variance *CV* and background variance *bg* of the assay and can thus be predicted without explicitly running experimental samples for every likely concentration. To predict the ROQ for a given set of affinity reagents and expected analyte concentrations, we can employ a three-step procedure. First, we use Equation 2 to calculate the expected measurement signal for 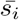 given analyte concentrations. This makes the implicit assumption that our simulated measurements are centered around the true noise-free readout. Next, we estimate the sample noise standard deviation *SD*_*i*_ from Equation 7. This can be established with experimental data by estimating the noise standard deviation from binding curve measurement replicates. Signal noise is partly a product of *bg*, but is also expected to increase relative to the signal magnitude with a given *CV*. Accordingly, we have updated *SD*_*i*_ to be a function of *CV* and *bg* (see Methods 1.6 for more detail). These variables can be determined either by literature search or empirically using binding curve data.

Finally, we apply our model to simulated values of 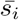 and 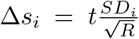 to calculate the upper and lower bounds for every analyte-affinity reagent pair, which we then convert to the estimated ROQ. We define the ROQ as the log_10_-difference between the upper and lower bounds of the quantification range: ROQ = log_10_(upper bound) − log_10_(lower bound), such that an increase of ROQ by one translates to a quantification range that is wider by one order of magnitude. The numerical value of the ROQ that is useful depends on the use case. However, for general intuition, a ROQ of 0.5 indicates that the true value (if the range encompasses it) lies within a 0.5 order of magnitude window, which is generally accepted to be good quantification. Conversely, a ROQ of three indicates a window size of three orders of magnitude, which is typically not useful in the context of physiological concentrations of biological molecules.

As a visual aid, we can plot the predicted ROQ as a heatmap for each intended analyte. Using this method, we simulated ROQ heatmaps for kyn in our example system (Fig. 3a), where *CV* and *bg* were estimated from the normalized binding curve and analyte-free control experimental data (*CV* = 0.053, *bg* = 8.7·10^−5^). Next, we compared how well our predicted quantitative performance matches real world measurements. As we observed experimentally, samples with lower concentrations of xa in the experiments presented in Figure 2d-f had a low ROQ for kyn, indicating superior quantitative precision, while samples with higher concentrations of xa (as in Fig. 2g-h) were predicted to have high ROQ (Fig. 3a, red crosses). To more extensively validate this heatmap, we also experimentally tested a broader range of samples with conditions including extremely low [xa] and [kyn], high [xa] and low [kyn], or higher [kyn] than was previously tested (Fig. 3a, red circles). The overlapping signal curves from our two affinity reagents for each of these additional experiments are shown in Figure 3b–g, and the respective lower and upper bounds of quantification are shown in Figure 3h and i for our model and the naïve model, respectively.

**Figure 3.**
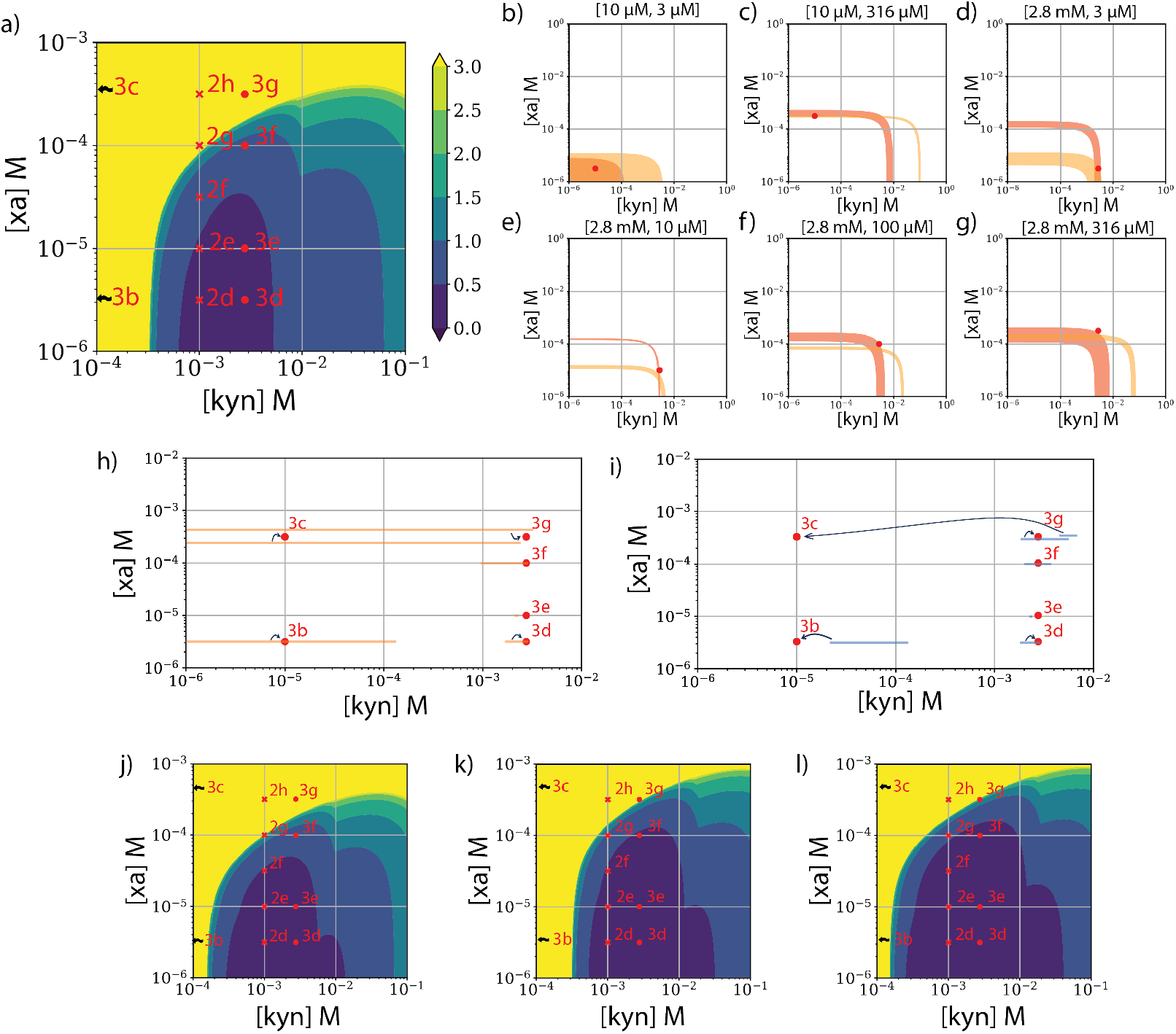
Using range of quantitation (ROQ) to predict assay confidence intervals. **a)** Discretized ROQ heatmap for kyn in an assay with SK1 and XA1, with an estimated CV of 0.2 and background of 0.01. Labels for each red cross or circle indicate figure panel number for their respective L-curves. Samples 3b and 3c are off the grid at 0.01 mM kyn, and have a ROQ of *>*3. **b–g)** Overlapping confidence interval curves from SK1 (red) and XA1 (orange) for various mixtures of kyn and xa, with actual analyte concentrations indicated by a red circle. **h-i)** High-confidence concentration ranges predicted for kyn using h, our method or i, the naïve assumption of specificity. Actual concentrations are marked with red circles. Arrows indicate which condition produces a given bar range when xa concentrations are the same and would otherwise produce overlapping bars. Exact values presented in SI Table 5. **j–l**) Revised ROQ heatmaps simulating conditions in which **j**, the background or **k**, the CV are reduced by 50%, or **l** in which the number of replicates used increases from three to five.

For samples with very low concentrations of kyn (Fig. 3b-c), our model predicts a large ROQ above 3, which is accurately reflected experimentally by the very large confidence intervals (Fig. 3h). This reflects the fact that background noise overwhelms the signal at such low concentrations. For samples with low concentrations of xa, and higher concentrations of kyn (Fig. 3d), our model predicts a ROQ *<* 0.5 for kyn. Our experimentally-derived confidence interval again confirms this, with good quantitative precision (Fig. 3h). Likewise, with high [kyn] and slightly higher [xa] (Fig. 3e), our experiment confirmed the model’s prediction that we would be able to quantify kyn with ROQ *<* 0.5, yielding a very narrow confidence interval that is slightly off from the true concentration. We note however, that the %CV 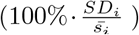 in the replicate SK1 signals for the mixture shown in Figure 3e is more than one order of magnitude lower than the average %CV for all mixed samples (2.8% vs 0.1%). This discrepancy suggests the possibility of measurement error leading to failure to accurately capture the true concentration. At higher concentrations of xa (Fig. 3f), our model predicted an ROQ between 0.5 and 1.0, which is reflected in the slightly wider experimentally derived ROQ. Accordingly, as shown in the sample condition in Figure 3g, further increasing [xa] was expected to significantly reduce the sensitivity of the signal output of SK1 to kyn and thus has a predicted poor ROQ *>* 3 — a result we confirmed experimentally and described in the previous section. Importantly, all of the experimentally calculated concentration ranges still encompass the true concentration even when the expected ROQ is broad, with the exception of the condition in Figure 3e (which, as explained above, is likely attributable to experimental error). In contrast, the naïve model generally produces deceptively narrow ROQs, but these only encompassed the true concentration in four out of eleven samples (Fig. 2i, 3i), all of which are for cases where cross-reactivity has a much lower effect (low [xa] and medium-high [kyn]).

These results show that our ROQ heatmaps accurately predict the experimental results in terms of quantitative precision, and should thus offer a robust tool for informing design decisions for assays. For example, our model can help in determining the maximum allowable noise that an assay can have to deliver reasonable precision at the analyte concentration range of interest. The ROQ heatmap in Figure 3a shows that a sample with 2.8 mM kyn and 100 µM xa (the condition shown in Fig. 3f) falls at the edge of zones describing ROQ of 1 and 1.5 orders of magnitude. If the goal is to bring the ROQ below *<* 0.5 for this analyte concentration, we can use our model to estimate how much the CV or background needs to be decreased to achieve this. As an exercise, we have simulated how the ROQ heatmap shifts when the background variance is lowered by a factor of four (Fig. 3j), when the coefficient of variance CV (Fig. 3k) is lowered by a factor of two, or even when the number of replicates used are increased from three to five (Fig. 3l). These results show that reducing the background standard deviation by 50% minimally changes the predicted ROQ for the sample with 2.8 mM kyn and 100 µM xa, whereas lowering the CV by 50% shifts the ROQ to the desired range of 0.5. This suggests that assay design in this case should prioritize the identification of opportunities to reduce the CV — for example, by implementing more stringent washing or referencing steps, depending on the assay [4]. Conversely, by keeping the assay the same, but running more sample replicates, drastically improves the ROQ for the same sample.

## Conclusion

In this work, we present a mathematical framework to enable the use of cross-reactive reagents for molecular quantification. We achieve this by using multiple affinity reagents with distinct analyte-affinity profiles alongside a generalized equilibrium-based model. For robustness, we also incorporated noise profiles into the framework to output a range of analyte concentrations that consistently encompass the true analyte concentration regardless of the concentration of interferent analytes.

We validated our theory with a proof-of-concept experiment focusing on quantifying kyn, a molecule for which no specific affinity reagents are available. Our framework provided tight quantification ranges that consistently contained the true kyn concentration under a range of conditions with varying concentrations of the cross-reactive analyte xa. Notably, even when the concentration of xa is so high that sensitivity drops significantly, the calculated range still encompassed the true concentration. Though a very large range was calculated in these saturating cases, consistently incorporating the true concentration is drastically more important in molecular measurement than incorrectly predicting a tight range that does not cover the true concentration. This furthermore highlights the importance of providing a range of concentrations rather than a single value for quantification. In contrast, a conventional Langmuir-based model that does not account for cross-reactivity consistently overestimated the kyn concentration, even as it generated a deceptively narrow range of quantification using our noise estimates. We have also demonstrated how to derive the ROQ metric for a given assay based on a limited set of experimental data, which enables the design of molecular detection assays that are better optimized for analyte quantification within an expected concentration range. Finally, to scale the framework up beyond the examples shown in this work, our solutions are provided as generalized solutions for an arbitrary number of analytes and reagents, and we have provided potential methods for handling different cross-reactivity scenarios in an experimentally tractable manner (SI Section 2.1).

Our framework helps turn the liability of cross-reactivity into an asset for achieving accurate quantification of analytes even with sub-optimal affinity reagents. This robust schema yields well-defined bounds of quantification that make it easier to assess the quality of assay results and determine conditions under which assay performance is likely to break down (even in cases of guaranteed specificity). To aid in scaling up to more reagents and analytes, our framework could be revised to resolve analyte concentrations in more complex, non-Langmuir binding models—for example, affinity reagents with multiple binding sites or scenarios in which the affinity reagent or analyte concentration is no longer guaranteed to be in excess of the other. To achieve this, the assays could be first run to generate the necessary data to understand the relevant binding interactions. If the binding interactions break assumptions stated in the methods, the generalized binding equation can be updated to resolve such physical changes. Furthermore, if non-linearities are introduced to the binding equations, our specific method of solving the equations with linear programming could be changed out to use other optimization solutions such as Bayesian estimators and more common optimization techniques like maximum likelihood estimation. Furthermore, integrating this framework with affinity reagent selection processes could potentially provide affinity reagent sets with varying degrees of cross-reactivity that improve the quantification of analytes in their expected concentration ranges, thereby minimizing the burden on affinity reagent selections.

In conclusion, this work provides a proof-of-concept framework for re-purposing the many existing cross-reactive affinity reagents to develop molecular detection assays that can achieve greater quantitative accuracy in complex samples, and thus could inform a reassessment of what constitutes a ‘successful’ affinity reagent selection process.

## 1 Methods

### 1.1 Generalized Model Derivation and Assumptions for Cross-Reactive Affinity Reagents

Our mathematical framework is applicable for any number of reagents (*m*) and analytes (*n*), although we highly suggest keeping *m* ≥ *n* to avoid an under-determined system of equations and minimize the ROQ. Here, we provide a generalized model and its derivation for the signal output *s*_*i*_ from a cross-reactive affinity reagent *A*_*i*_ (*i* ranges from 1 to *m*), which can bind to *n* analytes *T*_*j*_ (*j* ranges from 1 to *n*). Each of these *n* interactions has a respective association constant of 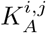:

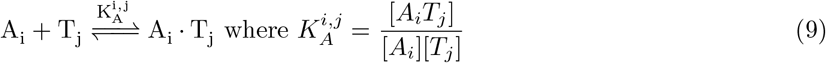

At equilibrium, the fraction of analyte *T*_*j*_ bound to *A*_*i*_ is a function of all interactions of *A*_*i*_ with all *n* analytes. To simplify all the possible binding dynamics, we assume the following: *A*_*i*_ can only bind to a single analyte at a time, analyte concentrations [*T*_*j*_] are significantly greater than the sum of all affinity reagent concentrations [*A*_*i*_], and there is only a single 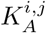 for each *A*_*i*_ binding to *T*_*j*_ [9, 10]. As such, the fraction of *A*_*i*_ bound to *T*_*j*_ is derived as follows:

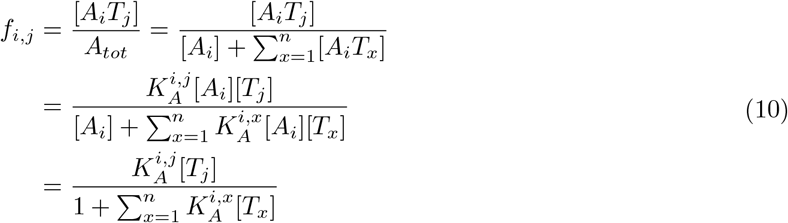

A normalized signal output from *A*_*i*_ will be a sum of fractions of *A*_*i*_ bound to *T*_*j*_:

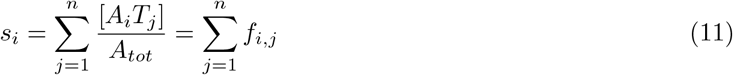

Combining equations 10 and 11 yields the following:

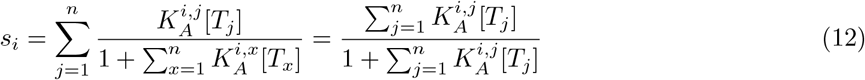

which describes the total signal output for *A*_*i*_ in a solution of *n* analytes. Note that when we assume there is only one analyte (*n* = 1), Equation 12 reduces to the Langmuir isotherm [11]. As we already assume that [ *T*_*j*_] *>>* [ *A*_*i*_], we also assume that the interactions of *m* other affinity reagents negligibly reduce the free *T*_*j*_ in solution, and thus any other affinity reagent *A*_*i*_ will follow the same equation as Equation 12.

### 1.2 Solving for the Upper and Lower Bounds of Concentrations

To calculate lower and upper bounds on all analyte concentrations, we assume that we have access to reliable upper and lower bounds for every signal readout *s*_*i*_. Our approach is a formalization of the visually intuitive approach of Fig. 1, where each affinity reagent’s signal *s*_*i*_ defines a set of analyte concentration combinations that could feasibly generate that signal. Overlapping these feasible sets thus defines the solution set of analyte concentrations that can explain all measurements. To determine this solution set mathematically, we begin by formally describing it using linear constraints (inequalities of the format 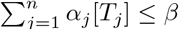 with some values *α*_*j*_ and *β*). We then determine the solution set’s extremes (i.e., lower and upper bounds for each analyte concentration) using off-the-shelf convex optimization solvers. These solvers have the advantage that they guarantee convergence to the same optimal solution.

Convex optimization solvers work on convex sets (i.e., contiguous shapes without any dents). The solution set forms such a convex set when plotted on a linear scale rather than a logarithmic one. To formally construct the solution set, we reshape all of our constraints into a linear format to yield linear constraints on the analyte concentrations and then sequentially apply them. These linear constraints can be thought of as cutting the space along a plane, and removing everything that is on the wrong side of the plane.

We begin constructing our convex set using the solubility limits 0 ≤ [*T*_*j*_] ≤ *T*_*max,j*_. The lower limits of 0 ≤ [*T*_*j*_] cut the space along the zero-plane of every dimension, leaving only the positive concentrations. Adding the upper solubility limits [*T*_*j*_] ≤*T*_*max,j*_ turns the space of feasible concentrations into a (hyper)rectangle.

Next, we convert the upper and lower bounds on the readout *s*_*i*_ into linear constraints on [*T*_*j*_]. For notational brevity, we rewrite Eq. 7 as

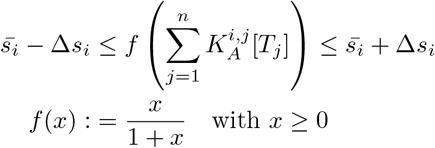

Because *f* is monotonically increasing for positive *x*, there exists an inverse function *f* ^−1^, which we can utilize to gain linear constraints on the analyte concentrations:

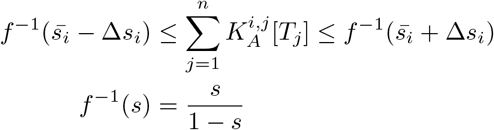

As an aside, if *f* is monotonically decreasing (e.g., for a signal-off binder), the inequalities change to be:

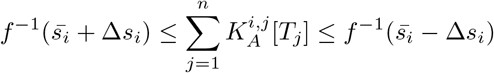

We can now cut our feasible space based on these linear constraints. Whereas the solubility limits were parallel with the axes, these linear constraints are more diagonal. In summary, our convex shape is described by the following constraints:

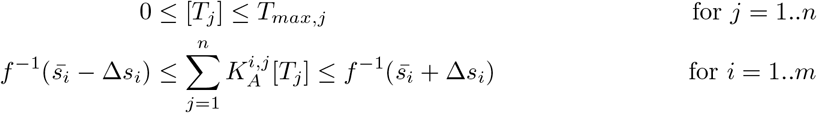

Having described the convex shape, we can query its properties using a convex optimization solver, such as CVXPY [13, 14]. The properties we care about for the purposes of this paper are the minimum and maximum values of each analyte concentration within the solution set. To find these values for a given analyte *T*_*k*_, we solve the following convex problems^3^:

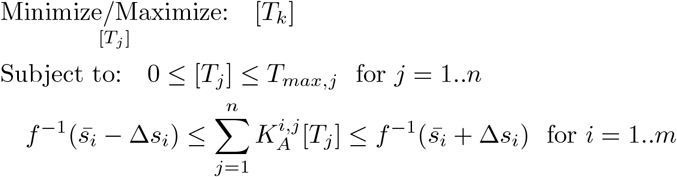

Note: To solve for ranges of quantification using the conventional naïve Langmuir model (as used for comparison in the experimental results) we pretended every analyte had a mono-specific affinity reagent. We set the *K*_*A*_ values of SK1 for xa and of XA1 for kyn to zero, and then used the same methods.

### 1.3 Reagents

DNA aptamers and displacement strands were chemically synthesized by Integrated DNA Technologies (IDT) with high-performance liquid chromatography (HPLC) purification (see SI Table 1 for sequences). L-kynurenine (K8625) and xanthurenic acid (D120804) were ordered from Sigma-Aldrich.

### 1.4 Assay Characterization via Plate Reader

Aptamers and displacement strands were developed and characterized in previously published work [7]. These affinity reagents were selected to undergo a conformational change upon binding to an analyte. Thus, when functionalized with fluorophore-quencher pairs, such binding events produce a fluorescence increase that can be read in a conventional plate-reader-based assay. We followed the same plate-reader-based assay with minor modifications. For initial stock solutions, analytes were dissolved in selection buffer (20 mM TrisHCl, pH 7.0, 30 mM NaCl, 5 mM KCl, 1 mM MgCl2, and 0.01% tween-20) at a concentration of 5 mM for kyn or 2 mM for xa. These were then serially diluted down to 0.123 µM kyn or 0.013 µM xa for binding curve concentrations. Master mixes of aptamer and displacement strand were created as a 10x solution of 500 nM aptamer and 2 or 1.25 µM displacement strand for XA1 and SK1, respectively. These mixtures were then pre-annealed in 100 µl aliquots by holding at 95ºC for five minutes and dropping by 1ºC every 30 seconds until reaching 4ºC in a Mastercycler X50 (Eppendorf).

To create the sample mixtures, 6 µl of 10x annealed master mix was combined with 54 ul of 1.1x concentrated target in 96 well, semi-skirted, LoBind twin.tec PCR plates (Eppendorf) for a total volume of 60 µL at 1x master mix and 1x target concentration. No-analyte controls were set up by adding 1x selection buffer instead of analytes. After sealing the plate, the samples were briefly vortexed and incubated on a rotator for 30 minutes in the dark at room temperature. The plate was then spun down at 1,000 rcf for 5 seconds. 50 µl of each sample were transferred to Corning 96-well half-area black-bottom polystyrene microplates (Thermo Fisher Scientific) for measurements at 25ºC on a Synergy H1 microplate reader (BioTek), with a filter cube (emission: 590/35, excitation: 538/63, gain: 55).

### 1.5 3-Parameter-Logistic (3-PL) Curve Fitting

All of our methods require specific values of 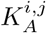, and require our signals *s*_*i*_ to be normalized to the range [0; 1]. We assume our unnormalized signals *u*_*i*_ follow the following 3-PL equation (Which is the same as 4-PL, but with a Hill coefficient of 1 since we assume one-to-one binding):

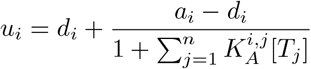

where *d*_*i*_ is the readout value from our assay when the affinity reagents are saturated, and *a*_*i*_ is the readout when analyte concentrations are zero. Given these parameters, we normalize our reads as follows:

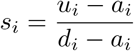

To determine *a*_*i*_, *d*_*i*_, and 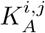, we measure the signal of each affinity reagent against varying concentrations of each analyte in isolation, i.e., the concentrations for all but one target analyte are zero. The equation that models this system simplifies to be:

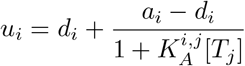

We then apply a least-squares fit to these parameters using the `curve fit` function of scipy [16]. Specifically, we solve the following optimization problem:

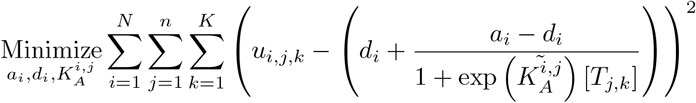

where *i* is the index for the affinity reagents, *j* is the index for the target analytes, and *k* is the index for the samples and replicates. We substitute 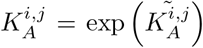 to make the optimization problem more stable. To ensure convergence, we also defined lower and upper bounds on all fitted parameters. The values of these bounds were chosen to be very lenient by being unrealistically small/large values (e.g., beyond detection limits of the detectors, and beyond solubility limits). Specifically, we set 0 ≤ *u*_*i*_, *d*_*i*_ ≤ 10^6^, and 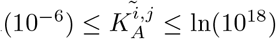(equating to limits of 1 aM and 1 MM on 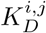).

When running these experiments for xa and kyn, we noticed that the highest concentrations from both analytes’ binding curves were producing uncharacteristically low readout values. This may have been due to potential quenching of fluorescence by the analytes at high concentrations [7]. For this reason, we ignored these highest concentrations to get more reliable fits. Full data are provided in the supplemental data file.

### 1.6 Calculation of assay error profile and background signal

To estimate the standard deviations we would expect from a given measurement, we assume that there are two sources of variance in our measurements. First, a constant background variance *bg*, second, a variance that scales with signal output according to a coefficient of variance 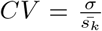. We then assume that these sources of variance are independent, such that the variance of a signal *s*_*k*_ is simply the sum of both sources of variance:

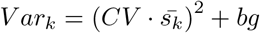

To estimate these two parameters *bg* and *CV*, we first normalized the signals from the binding curve/analyte-free controls as described above in the 3-PL curve fitting methods section. For each sample, we then calculated the average 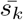 and variance *V ar*_*k*_ based on triplicate measurements. Finally, we determined *bg* and *CV* by solving the following optimization problem:

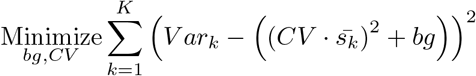

where *k* is the index for the samples. For a given signal *s*_*i*_, we can now estimate the sample standard deviation

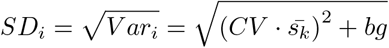

and the Δ*s*_*i*_ from Equation 7:

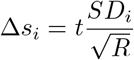

where *R* is the number of measurement replicates we are simulating, and *t* is the 97.5th percentile of the t-distribution with (*R* − 1) degrees of freedom.

### 1.7 Statistics and Reproducibility

All samples were measured in triplicate. All attempts are presented in the figure plots except for the highest concentration samples in the binding curve due to potential quenching of fluorescence by the analytes (discussed in Methods 1.5). These data are kept in the source dataset. No statistical method was used to predetermine sample size. The investigators were not blinded to allocation during experiments and outcome assessment.

## 1.8 Data Availability

The source data generated in this study are provided with this paper.

## 1.9 Code Availability

Code is available here: https://github.com/sohlab/cr_quant

## 1.10 Acknowledgements

S.S.N acknowledges support from SGF (Stanford Graduate Fellowship in Science and Engineering) and the NSF Graduate Research Fellowship Program (GRFP). We acknowledge Brandon Wilson for working with S.S.N in developing the first version of the generalized binding equation with cross-reactivity. We also thank Leighton Wan for providing guidance on using his and Alex Yoshikawa’s XA-1 and SK-1 aptamers. Thank you also to Benjamin Wollant and Nicholas Vitale for their insightful comments in our manuscript. We would like to thank Mr. Michael Eisenstein for his assistance in editing of the manuscript.

## 1.11 Author Contributions

S.S.N. ideation, formal problem statement, experimental set-up, code support, and writing the paper; L.A.H. overlapping solution idea, main code implementation, and writing the paper; A.M.A. experimental validation; H.T.S. general support, formal problem statement, and writing the paper.

## 2 Supplemental Information

### 2.1 Strategies for addressing different categories of cross-reactive analytes

Effectively handling the large number of analytes that might bind to an affinity reagent requires us to first identify which ones have a substantial impact on the signal output. Such impacts depend on the variability of the cross-reactive analyte concentration and the ratio of the effective signal contribution of the cross-reactive analyte to intended analyte 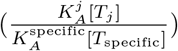. To tackle this, we suggest categorizing cross-reactive analytes into three groups: “high”, “low”, and “constant” cross-reactivity. Subsequently, we apply our model to each group with decreasing levels of detail.

The most important cross-reactive analytes are those in the “high” cross-reactivity category, which are expected to vary a lot in a sample and for which the ratio of cross-reactivity to specific signal is high (⪆ 1). These “high” analytes can typically be identified from the literature for the affinity reagent *A*_*i*_ and by understanding analyte-relevant molecular pathways, and should be individually modeled. It is up to the researcher’s discretion how much error they can tolerate. The impact of xa on kyn quantification is an example of such a high cross-reactivity analyte, since xa is expected to vary a lot in physiological conditions and the ratio of cross-reactivity in these conditions with the available affinity reagents for xa and kyn is high.

Low cross-reactivity analytes are those for which the ratio of cross-reactivity to specific signal is low, but [*T*_*j*_] can still vary. It might not be worth extracting out *K*_*A*_ values for each analyte in this category. However, we suggest approximating these into a total combined association constant 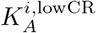. We can approximate this by making dilutions of the sample matrix (without *T*_specific_) and measuring the change in signal output from each affinity reagent.

Finally, constant cross-reactive analytes are those which are known to cross-react, but their concentration is expected to stay relatively constant. These analytes contribute to the background signal, causing a constant shift upwards in the binding curve. Depending on how much the cross-reactivity to specific signal ratio changes, there could be a constant reduction in the dynamic range of the signal. This does not introduce bias into quantification, and thus is beyond the scope of our current study. For more information about reducing non-specific binding, interested readers can explore other techniques mentioned in the introduction. For constant cross-reactivity analytes, one should always be careful to verify the assumption of the signal contribution being constant. Wrongfully dismissing off-target binding as “constant” cross-reactivity will have the same effect as not accounting for cross-reactivity at all, as in the naive methods discussed in this paper.

### 2.2 An analytical solution to the cross-reactivity model leads to unstable solutions

In the case of no noise, we can solve for the analyte concentrations analytically:

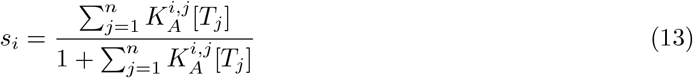

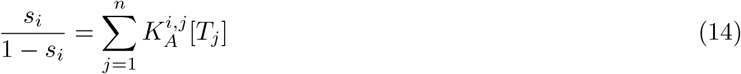

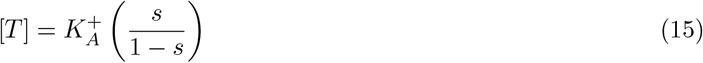

where [*T*] is a vector of analyte concentrations, 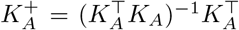 is the pseudo-inverse of the matrix of association constants *K*_*A*_, and *s* is a vector of the signal readouts from all affinity reagents. We see that this system can only be solved if there is no saturated affinity reagent readout with *s*_*i*_ = 1, and if the matrix 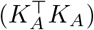is invertible, i.e., we have enough affinity reagents whose association constants are linearly independent from one another.

While this set of equations is easily solvable, it runs into the problem of being sensitive to noise without giving the user any feedback for when they may trust the readouts. To illustrate this problem, we will analyze the case of two analytes and two affinity reagents. The closed-form solution of the above equations are:

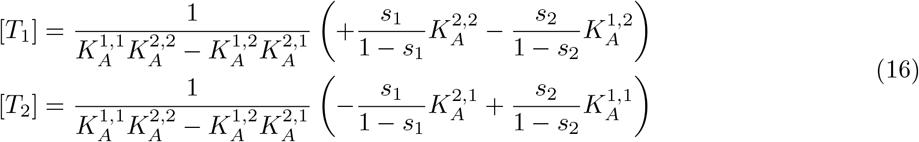

Looking at the example case where

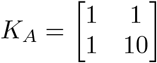

with [*T*_1_] = 1, [*T*_2_] = 0, we get the signal readouts *s*_1_ = *s*_2_ = 0.5. Using the above equations, we can now observe the effect of small errors in *s*_2_. For *s*_2_ = 0.5 + 10^−6^, we get [*T*_2_] = 4 · 10^−8^. For *s*_2_ = 0.5 + 10^−3^, we get [*T*_2_] = 5 · 10^−5^, a change of three orders in magnitude. This shows that even small errors in *s*_2_ can change this method’s estimate of [*T*_2_] by orders of magnitude, without alerting the user to this sensitivity to noise.

In contrast, using our method gives an intuitive understanding of what is happening in this scenario: This example’s analyte concentrations are in a regime of high accuracy for [*T*_1_] and low accuracy for [*T*_2_], which is due to the values 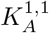 and 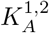 being too close, as is reflected in the corresponding ROQ heatmap. Our method reports this by only yielding an upper bound for [*T*_2_], instead of attempting to make an accurate estimate. For this reason, we argue that the presentation of concentrations as feasible ranges on a logarithmic scale rather than single-number estimates can prevent misunderstandings when considering assay results.

### Supplementary Figures

**Figure 4.**
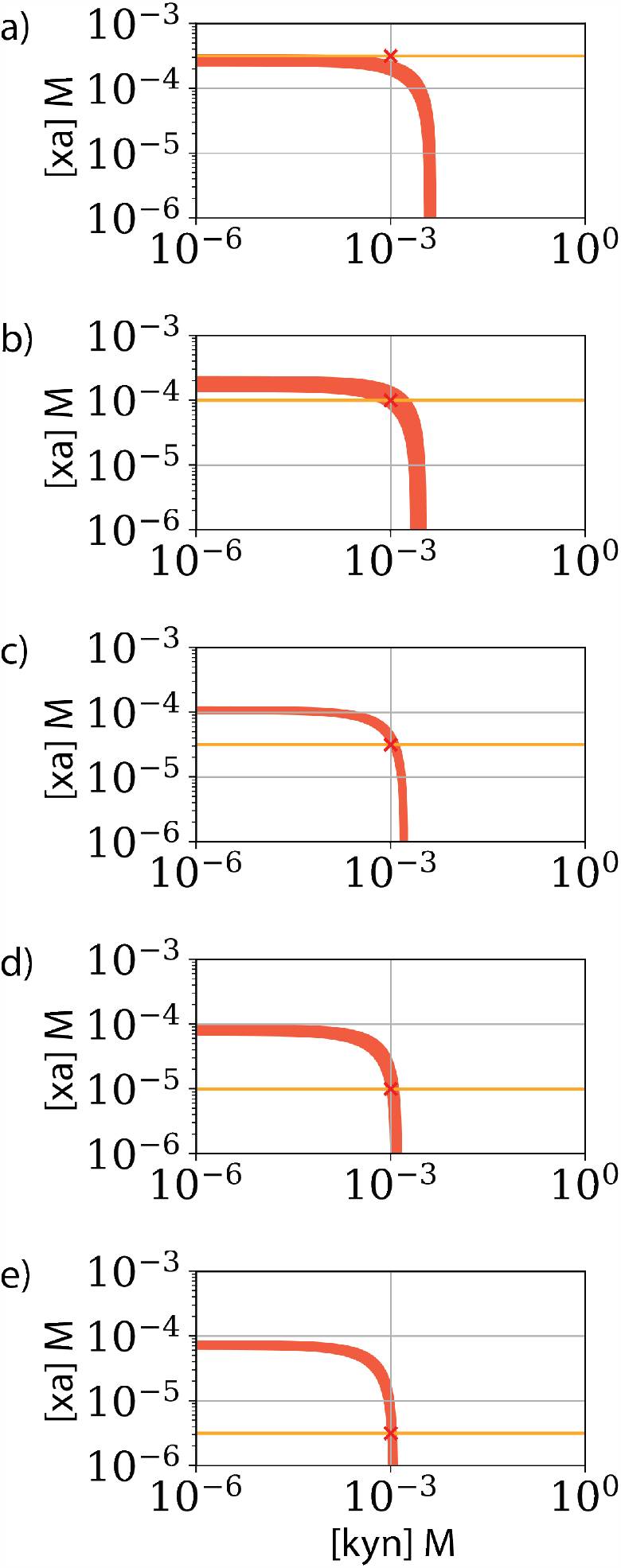
Demonstration of inherent limitations of affinity reagent SK1. For the experimental conditions and resultant signals of 1mM kyn and increasing xa from 3 µM to 316 µM **a-e)** as discussed in the main section. Even when overlapping measurements from SK1 (red L-shapes) with perfect information about the true xa concentration (yellow horizontal line), the overlapped region may not be bounded for kyn. This occurs when xa concentrations approach the 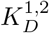 of SK1 (in this case 0.13 mM) which is about the conditions shown **d-e**. At these points, the xa line overlaps with the lower end of SK1 signal output. Another way of viewing this is that as the xa concentration increases, more SK1 affinity reagents are occupied by xa, leaving fewer affinity reagents to interact with kyn. This causes the ROQ for kyn to increase in size for high concentrations of xa.

## Supplementary Tables

**Table 1:**
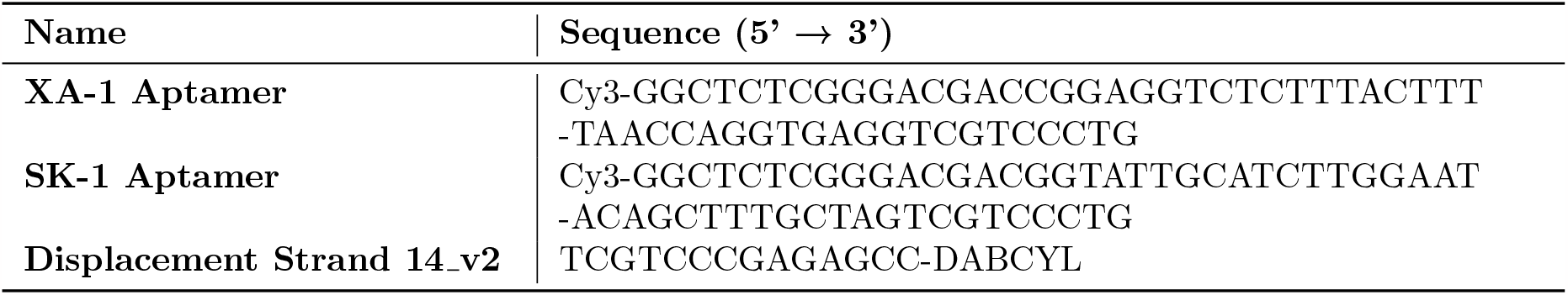
DNA aptamer and displacement strands sequence.

**Table 2:**
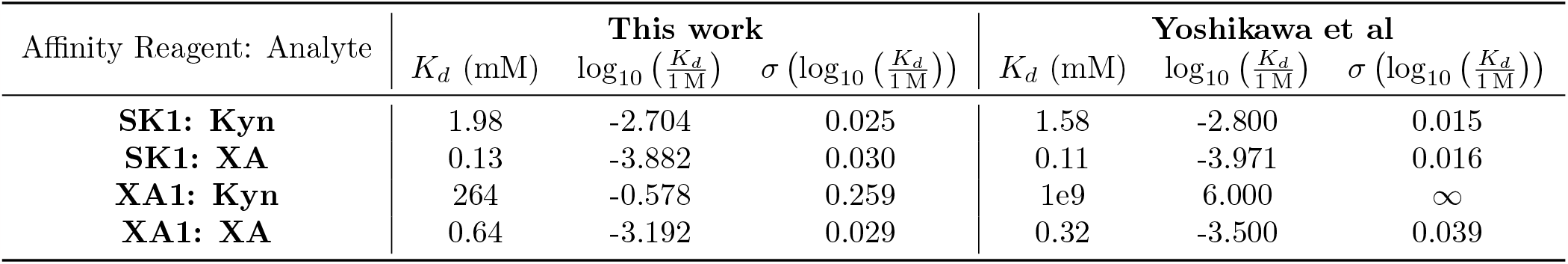
*K*_*d*_ fits for this work’s binding curve data versus the original work by Yoshikawa et al. [7] when fit with a 3-parameter logistic curve. The values for XA1: Kyn from the original work by Yoshikawa et al. of *K*_*d*_ = 10^6^M and 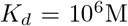 are due to the fitting algorithm hitting the upper bound for 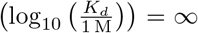, which we defined in section 1.5 to ensure convergence.

| Affinity Reagent: Analyte | This work |  |  | Yoshikawa et al |  |  |
| --- | --- | --- | --- | --- | --- | --- |
| | $K_d$ (mM) | $\log_{10}(\frac{K_d}{1\text{M}})$ | $\sigma(\log_{10}(\frac{K_d}{1\text{M}}))$ | $K_d$ (mM) | $\log_{10}(\frac{K_d}{1\text{M}})$ | $\sigma(\log_{10}(\frac{K_d}{1\text{M}}))$ |
| **SK1: Kyn** | 1.98 | -2.704 | 0.025 | 1.58 | -2.800 | 0.015 |
| **SK1: XA** | 0.13 | -3.882 | 0.030 | 0.11 | -3.971 | 0.016 |
| **XA1: Kyn** | 264 | -0.578 | 0.259 | 1e9 | 6.000 | ∞ |
| **XA1: XA** | 0.64 | -3.192 | 0.029 | 0.32 | -3.500 | 0.039 |

**Table 3:**
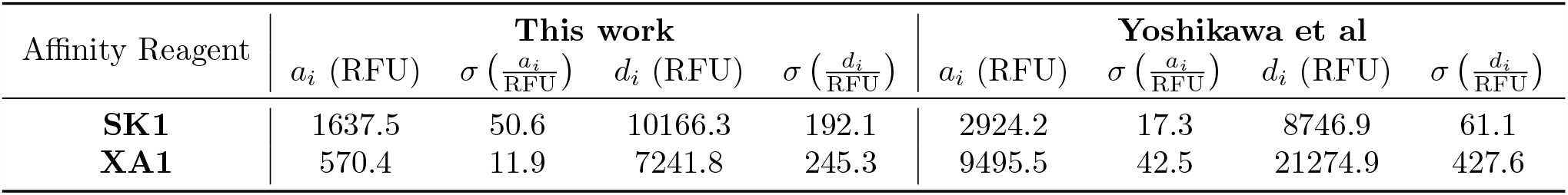
Lower bound values *a*_*i*_ and upper bound values *d*_*i*_ from this work’s binding curve data versus the original work by Yoshikawa et al. [7] when fit with a 3-parameter logistic curve. Note that we assumed the change in fluorescence to be identical no matter which analyte binds to an affinity reagent. This means we only have a single value for *a*_*i*_ and *d*_*i*_ per affinity reagent, instead of one for each pairing of affinity reagent and analyte.

**Table 4:**
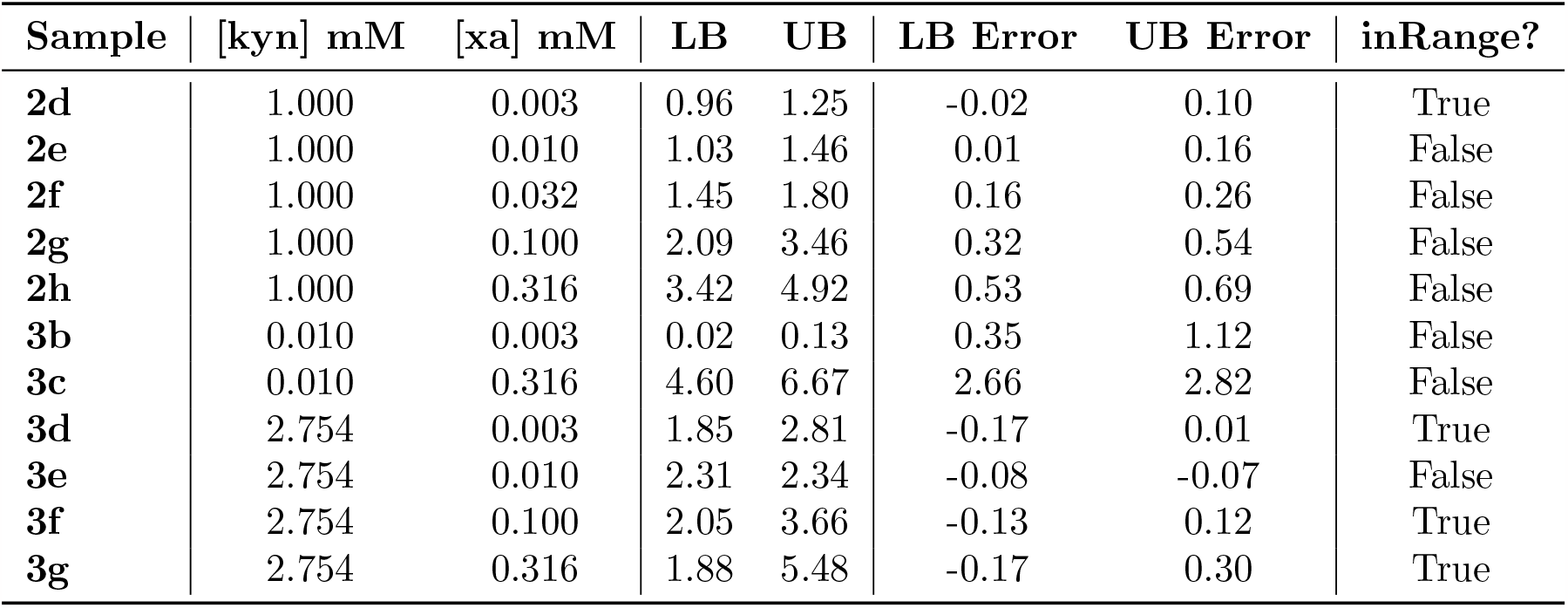
Upper and lower bounds (UB, LB) of kyn quantification in mM for the naive Langmuir model. Error calculated by log(LB or UB) - log([kyn]). ‘inRange’ indicates if estimated bounds hold the true kyn concentration. Sample names refer to figure panels in which curves from the relevant mixtures are presented.

**Table 5:**
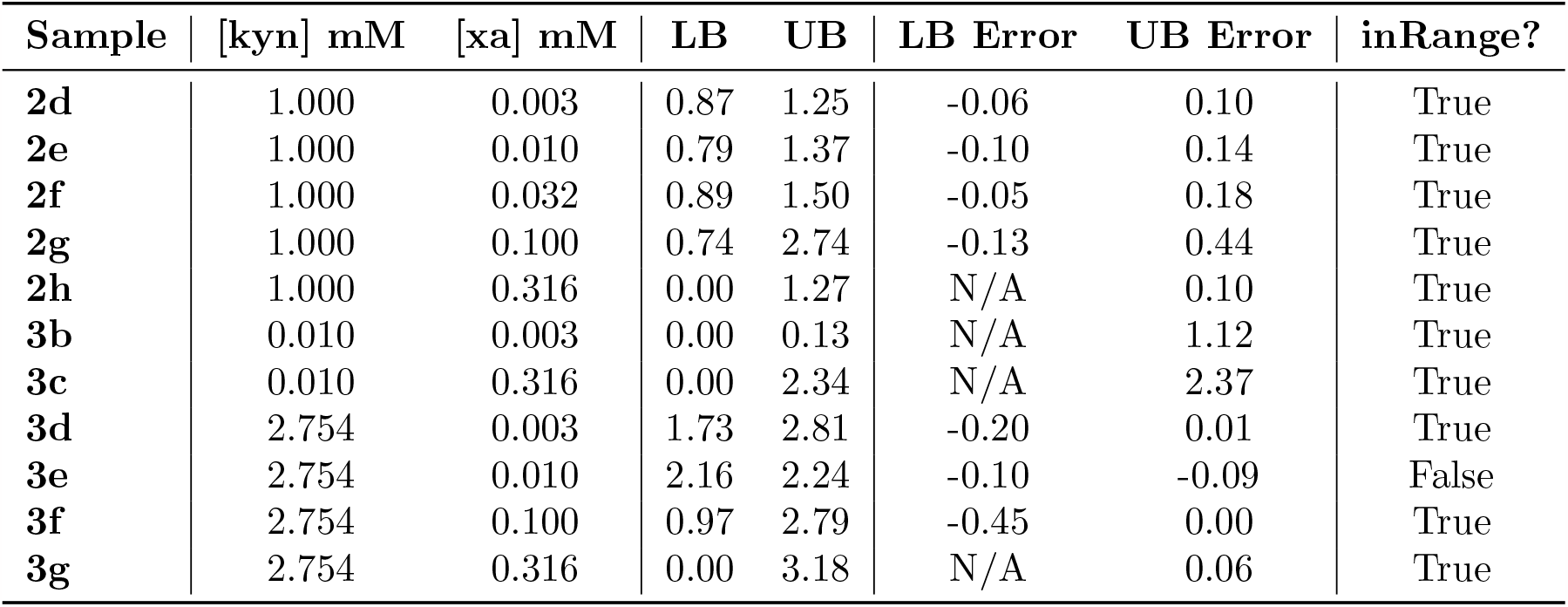
Upper and lower bounds (UB, LB) of kyn quantification in mM for our cross-reactivity model. Error calculated by log(LB or UB) - log([kyn]). N/A provided if bound is 0. ‘inRange’ indicates if estimated bounds hold the true kyn concentration. Sample names refer to figure panels in which curves from the relevant mixtures are presented.

## Footnotes

1 As the number of cross-reactive analytes increases, the exact handling of the analytes must change for experimental tractability. We provided potential methods to handle three cases of cross-reactivity—high, low and constant cross-reactivity—in SI Section 2.1.

2 As [xa] approaches the *K*_*D*_ of SK1 to xa 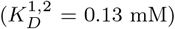, the sensitivity that SK1 has to [kyn] drastically drops. In SI Figure 4, we visually show that even with perfect knowledge of [xa] this drop in sensitivity forces the range of kyn quantification in this regime to be very large. This is also generally observed in the binding curves with increased *T*_2_ as previously depicted in Figure 1b.

3 Notation explanation: We use the notation by Boyd and Vandenberghe [15]. The indices are defined as follows: *k* is the fixed index of the target analyte whose bounds we are querying, *j* is a variable index over the analytes, *i* is a variable index over the readouts.

