## Supplemental Information for "Theoretical framework and experimental validation of multiplexed analyte quantification using cross-reactive affinity reagents"

#### 2.2 An analytical solution to the cross-reactivity model leads to unstable solutions

In the case of no noise, we can solve for the analyte concentrations analytically:

$$s_i = \frac{\sum_{j=1}^{n} K_A^{i,j}[T_j]}{1 + \sum_{j=1}^{n} K_A^{i,j}[T_j]} \tag{13}$$

$$\frac{s_i}{1 - s_i} = \sum_{j=1}^{n} K_A^{i,j}[T_j] \tag{14}$$

$$[T] = K_A^{+}\left(\frac{s}{1 - s}\right) \tag{15}$$

where $[T]$ is a vector of analyte concentrations, $K_A^{+} = (K_A^{\top}K_A)^{-1}K_A^{\top}$ is the pseudo-inverse of the matrix of association constants $K_A$, and $s$ is a vector of the signal readouts from all affinity reagents. We see that this system can only be solved if there is no saturated affinity reagent readout with $s_i = 1$, and if the matrix $(K_A^{\top}K_A)$ is invertible, i.e., we have enough affinity reagents whose association constants are linearly independent from one another.

While this set of equations is easily solvable, it runs into the problem of being sensitive to noise without giving the user any feedback for when they may trust the readouts. To illustrate this problem, we will analyze the case of two analytes and two affinity reagents. The closed-form solution of the above equations are:

$$
\begin{aligned}
[T_1] &= \frac{1}{K_A^{1,1}K_A^{2,2} - K_A^{1,2}K_A^{2,1}} \left( +\frac{s_1}{1-s_1}K_A^{2,2} - \frac{s_2}{1-s_2}K_A^{1,2} \right) \\
[T_2] &= \frac{1}{K_A^{1,1}K_A^{2,2} - K_A^{1,2}K_A^{2,1}} \left( -\frac{s_1}{1-s_1}K_A^{2,1} + \frac{s_2}{1-s_2}K_A^{1,1} \right)
\end{aligned} \tag{16}
$$

Looking at the example case where

$$
K_A = \begin{bmatrix} 1 & 1 \\ 1 & 10 \end{bmatrix}
$$

with $[T_1] = 1$, $[T_2] = 0$, we get the signal readouts $s_1 = s_2 = 0.5$. Using the above equations, we can now observe the effect of small errors in $s_2$. For $s_2 = 0.5 + 10^{-6}$, we get $[T_2] = 4 \cdot 10^{-8}$. For $s_2 = 0.5 + 10^{-3}$, we get $[T_2] = 5 \cdot 10^{-5}$, a change of three orders in magnitude. This shows that even small errors in $s_2$ can change this method's estimate of $[T_2]$ by orders of magnitude, without alerting the user to this sensitivity to noise.

In contrast, using our method gives an intuitive understanding of what is happening in this scenario: This example's analyte concentrations are in a regime of high accuracy for $[T_1]$ and low accuracy for $[T_2]$, which is due to the values $K_A^{1,1}$ and $K_A^{1,2}$ being too close, as is reflected in the corresponding ROQ heatmap. Our method reports this by only yielding an upper bound for $[T_2]$, instead of attempting to make an accurate estimate. For this reason, we argue that the presentation of concentrations as feasible ranges on a logarithmic scale rather than single-number estimates can prevent misunderstandings when considering assay results.

#### Supplementary Figures

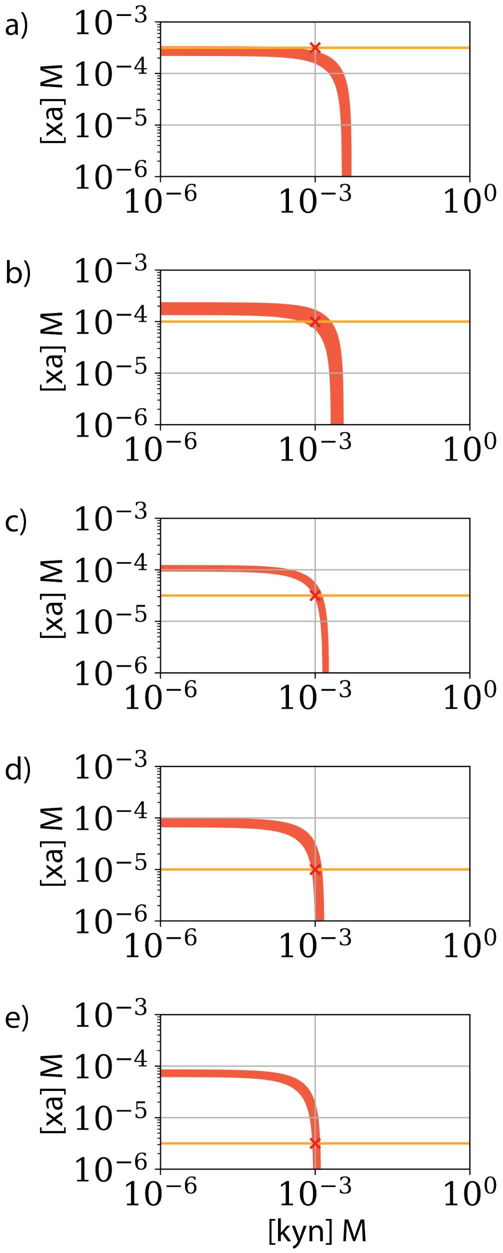


Figure 4: Demonstration of inherent limitations of affinity reagent SK1. For the experimental conditions and resultant signals of 1mM kyn and increasing xa from 3 µM to 316 µM **a-e)** as discussed in the main section. Even when overlapping measurements from SK1 (red L-shapes) with perfect information about the true xa concentration (yellow horizontal line), the overlapped region may not be bounded for kyn. This occurs when xa concentrations approach the $K_D^{1;2}$ of SK1 (in this case 0.13 mM) which is about the conditions shown **d-e**. At these points, the xa line overlaps with the lower end of SK1 signal output. Another way of viewing this is that as the xa concentration increases, more SK1 affinity reagents are occupied by xa, leaving fewer affinity reagents to interact with kyn. This causes the ROQ for kyn to increase in size for high concentrations of xa.

### Supplementary Tables

Table 1: DNA aptamer and displacement strands sequence

| Name | Sequence (5' → 3') |
|---|---|
| **XA-1 Aptamer** | Cy3-GGCTCTCGGGACGACCGGAGGTCTCTTTACTTT<br>-TAACCAGGTGAGGTCGTCCCTG |
| **SK-1 Aptamer** | Cy3-GGCTCTCGGGACGACGGTATTGCATCTTGGAAT<br>-ACAGCTTTGCTAGTCGTCCCTG |
| **Displacement Strand 14_v2** | TCGTCCCGAGAGCC-DABCYL |

Table 2: $K_d$ fits for this work's binding curve data versus the original work by Yoshikawa et al. [7] when fit with a 3-parameter logistic curve. The values for XA1: Kyn from the original work by Yoshikawa et al. of $K_d = 10^6$M and $\sigma\left(\log_{10}\left(\frac{K_d}{1\,\mathrm{M}}\right)\right) = \infty$ are due to the fitting algorithm hitting the upper bound for $K_A^{i,j}$, which we defined in section 1.5 to ensure convergence.

| Affinity Reagent: Analyte | **This work** | | | **Yoshikawa et al** | | |
|---|---|---|---|---|---|---|
| | $K_d$ (mM) | $\log_{10}\left(\frac{K_d}{1\,\mathrm{M}}\right)$ | $\sigma\left(\log_{10}\left(\frac{K_d}{1\,\mathrm{M}}\right)\right)$ | $K_d$ (mM) | $\log_{10}\left(\frac{K_d}{1\,\mathrm{M}}\right)$ | $\sigma\left(\log_{10}\left(\frac{K_d}{1\,\mathrm{M}}\right)\right)$ |
| **SK1: Kyn** | 1.98 | -2.704 | 0.025 | 1.58 | -2.800 | 0.015 |
| **SK1: XA** | 0.13 | -3.882 | 0.030 | 0.11 | -3.971 | 0.016 |
| **XA1: Kyn** | 264 | -0.578 | 0.259 | 1e9 | 6.000 | $\infty$ |
| **XA1: XA** | 0.64 | -3.192 | 0.029 | 0.32 | -3.500 | 0.039 |

Table 3: Lower bound values $a_i$ and upper bound values $d_i$ from this work's binding curve data versus the original work by Yoshikawa et al. [7] when fit with a 3-parameter logistic curve. Note that we assumed the change in fluorescence to be identical no matter which analyte binds to an affinity reagent. This means we only have a single value for $a_i$ and $d_i$ per affinity reagent, instead of one for each pairing of affinity reagent and analyte.

| Affinity Reagent | **This work** | | | | **Yoshikawa et al** | | | |
|---|---|---|---|---|---|---|---|---|
| | $a_i$ (RFU) | $\sigma\left(\frac{a_i}{\mathrm{RFU}}\right)$ | $d_i$ (RFU) | $\sigma\left(\frac{d_i}{\mathrm{RFU}}\right)$ | $a_i$ (RFU) | $\sigma\left(\frac{a_i}{\mathrm{RFU}}\right)$ | $d_i$ (RFU) | $\sigma\left(\frac{d_i}{\mathrm{RFU}}\right)$ |
| **SK1** | 1637.5 | 50.6 | 10166.3 | 192.1 | 2924.2 | 17.3 | 8746.9 | 61.1 |
| **XA1** | 570.4 | 11.9 | 7241.8 | 245.3 | 9495.5 | 42.5 | 21274.9 | 427.6 |

Table 4: Upper and lower bounds (UB, LB) of kyn quantification in mM for the naive Langmuir model. Error calculated by log(LB or UB) - log([kyn]). 'inRange' indicates if estimated bounds hold the true kyn concentration. Sample names refer to figure panels in which curves from the relevant mixtures are presented.

| Sample | [kyn] mM | [xa] mM | LB | UB | LB Error | UB Error | inRange? |
|---|---|---|---|---|---|---|---|
| **2d** | 1.000 | 0.003 | 0.96 | 1.25 | -0.02 | 0.10 | True |
| **2e** | 1.000 | 0.010 | 1.03 | 1.46 | 0.01 | 0.16 | False |
| **2f** | 1.000 | 0.032 | 1.45 | 1.80 | 0.16 | 0.26 | False |
| **2g** | 1.000 | 0.100 | 2.09 | 3.46 | 0.32 | 0.54 | False |
| **2h** | 1.000 | 0.316 | 3.42 | 4.92 | 0.53 | 0.69 | False |
| **3b** | 0.010 | 0.003 | 0.02 | 0.13 | 0.35 | 1.12 | False |
| **3c** | 0.010 | 0.316 | 4.60 | 6.67 | 2.66 | 2.82 | False |
| **3d** | 2.754 | 0.003 | 1.85 | 2.81 | -0.17 | 0.01 | True |
| **3e** | 2.754 | 0.010 | 2.31 | 2.34 | -0.08 | -0.07 | False |
| **3f** | 2.754 | 0.100 | 2.05 | 3.66 | -0.13 | 0.12 | True |
| **3g** | 2.754 | 0.316 | 1.88 | 5.48 | -0.17 | 0.30 | True |

Table 5: Upper and lower bounds (UB, LB) of kyn quantification in mM for our cross-reactivity model. Error calculated by log(LB or UB) - log([kyn]). N/A provided if bound is 0. 'inRange' indicates if estimated bounds hold the true kyn concentration. Sample names refer to figure panels in which curves from the relevant mixtures are presented.

| Sample | [kyn] mM | [xa] mM | LB | UB | LB Error | UB Error | inRange? |
|---|---|---|---|---|---|---|---|
| **2d** | 1.000 | 0.003 | 0.87 | 1.25 | -0.06 | 0.10 | True |
| **2e** | 1.000 | 0.010 | 0.79 | 1.37 | -0.10 | 0.14 | True |
| **2f** | 1.000 | 0.032 | 0.89 | 1.50 | -0.05 | 0.18 | True |
| **2g** | 1.000 | 0.100 | 0.74 | 2.74 | -0.13 | 0.44 | True |
| **2h** | 1.000 | 0.316 | 0.00 | 1.27 | N/A | 0.10 | True |
| **3b** | 0.010 | 0.003 | 0.00 | 0.13 | N/A | 1.12 | True |
| **3c** | 0.010 | 0.316 | 0.00 | 2.34 | N/A | 2.37 | True |
| **3d** | 2.754 | 0.003 | 1.73 | 2.81 | -0.20 | 0.01 | True |
| **3e** | 2.754 | 0.010 | 2.16 | 2.24 | -0.10 | -0.09 | False |
| **3f** | 2.754 | 0.100 | 0.97 | 2.79 | -0.45 | 0.00 | True |
| **3g** | 2.754 | 0.316 | 0.00 | 3.18 | N/A | 0.06 | True |
